# Multiple genetic loci affect place learning and memory performance in *Drosophila melanogaster*

**DOI:** 10.1101/580092

**Authors:** P.A. Williams-Simon, C. Posey, S. Mitchell, E. Ng’oma, J.A. Mrkvicka, T. Zars, E.G. King

## Abstract

Learning and memory are critical functions for all animals, giving individuals the ability to respond to changes in their environment. Within populations, individuals vary, however the mechanisms underlying this variation in performance are largely unknown. Thus, it remains to be determined what genetic factors cause an individual to have high learning ability, and what factors determine how well an individual will remember what they have learned. To genetically dissect learning and memory performance, we used the DSPR, a multiparent mapping resource in the model system *Drosophila melanogaster,* consisting of a large set of recombinant inbred lines (RILs) that naturally vary in these and other traits. Fruit flies can be trained in a “heat box” to learn to remain on one side of a chamber (place learning), and can remember this (place memory) over short timescales. Using this paradigm, we measured place learning and memory for ∼49,000 individual flies from over 700 DSPR RILs. We identified 16 different loci across the genome that significantly affect place learning and/or memory performance, with 5 of these loci affecting both traits. To identify transcriptomic differences associated with performance, we performed RNA-Seq on pooled samples of 7 high performing and 7 low performing RILs for both learning and memory and identified hundreds of genes with differences in expression in the two sets. Integrating our transcriptomic results with the mapping results allowed us to identify nine promising candidate genes, advancing our understanding of the genetic basis underlying natural variation in learning and memory performance.

## Introduction

The ability to learn and remember are critically important, allowing individuals to adjust their behavior in response to stimuli to cope with changing environments. Learning and memory performance vary widely in different contexts across the animal kingdom, with some species having evolved higher learning and/or memory abilities depending on the selective pressures they have been exposed to ^1–5^. Behavioral experiments in butterflies ^6, 7^, chickadees ^8, 9^, moths ^10, 11^, honey bees ^12–14^, fruit flies ^15, 16^, rodents ^17–19^, and humans ^20^ have shown collectively that there is genetically based variation in these traits and have established the different types of learning and memory within species. This variation provides the raw material for natural selection to act on when higher or lower learning and memory performance is selected for in a population. Experimental evolution studies have empirically demonstrated that populations can evolve higher or lower learning performance ^21–23^. These results show that there are natural genetic variants that give rise to differences in performance in learning and memory. The identity of the vast majority of these variants in nearly all populations, however, remains unknown (for exceptions see: ^24, 25^).

Many empirical studies aim to understand the mechanistic basis of different types of learning and memory within the broad context of animal behavior, using both classical and operant learning and memory paradigms. Specifically, many previous molecular genetic studies have identified individual genes involved in learning, and memory in many different animal models, such as *Caenorhabditis elegans (C. elegans)* ^26–28^, *Mus musculus* ^29–31^ and *Drosophila melanogaster (D. melanogaster)* ^32–36^. For example, in *C. elegans*, mutants with a defective *nmr-1* encoding an N-methyl-D-aspartate (NMDA) receptor subunit fail to form both short-term and long-term memories ^28^. In fruit flies, mutations in the *rutabaga* (*rut*) gene ^33, 35^ and in the *dunce* (*dnc*) gene ^32, 37^ have both been shown to influence multiple types of learning and memory. These studies, along with follow up studies showing the mechanism of action of these genes ^38–41^, have provided crucial information about which genes are required to function for proper learning and memory.

Despite this success, it is not clear whether segregating variants within these same genes are causing the variation observed in performance among individuals in natural populations. It is possible that the genes identified via these approaches are so central to the processes of learning and memory that their function is highly conserved, and most of the individual-level variation in learning and memory is due to genetic variants in other genes ^4^. Another reason to expect that the variants identified via mutant studies might not correspond to those seen in natural populations is that many of these mutants show deleterious pleiotropic effects. For example, severe *dnc* mutant flies have female sterility and partial lethality ^39, 42^, phenotypes that would be strongly selected against in the wild. Thus, identifying the natural genetic variants underlying why some individuals in a population perform better or worse than others is critical to our understanding of the mechanistic basis of variability in learning and memory among individuals.

Given the complex processes that govern learning and memory, which certainly involve a large number of potentially interacting genes, single gene approaches do not necessarily capture what happens on a systematic or organismal level ^4^. Mapping studies in natural populations, such as quantitative trait loci (QTL) studies and genome-wide association studies (GWAS), have the potential to allow for the identification of multiple genetic variants influencing a complex trait in the natural genetic context in which the variants occur. However, genetic mapping studies have been challenging to perform for learning and memory phenotypes. Assaying learning and memory is often labor-intensive, requiring repeated behavioral trials ^1^, making it difficult to assay the large numbers of individuals typically required for high power to detect all but the highest effect QTL ^43–45^. In human populations, while learning, memory, and general cognitive function are known to be heritable ^4, 20^, they are also influenced strongly by a suite of environmental factors, which obviously cannot be systematically controlled in human studies. High levels of variation in those environmental factors will negatively affect the power to detect a genetic association ^43–45^. There have been a few successes where a causative natural variant for learning or memory has been identified. These include a GWAS in humans of short-term working memory that identified a polymorphism in the *SCN1A* gene, a voltage-gated sodium channel ^25^, a study in fruit flies linking the *foraging* gene, a cGMP-dependent protein kinase, to associative olfactory learning and memory ^24^, and a QTL mapping study in mice implicating the *Hcn1* gene, a hyperpolarization-activated cyclic nucleotide-gated channel, in fear conditioning ^46^. However, by and large, very few QTL for learning and memory have been discovered (but see ^47–50^, and even fewer specific causative variants (aside from the few exceptions noted above) have been identified in any system thus far.

Multiparental populations, consisting of a large number of recombinant inbred lines (RILs) generated from multiple inbred founder lines crossed for multiple generations, have the potential to allow for high-resolution, highly powered genetic mapping studies. The *Drosophila* Synthetic Population Resource (DSPR) is one such multiparental population, consisting of two sets of ∼800 RILs, each generated from an 8-way, 50-generation cross ^44, 51, 52^. This system, with a large number of RILs and a long period of crossing allows for highly powered QTL mapping to an interval that typically includes tens of genes rather than the wide intervals including hundreds of genes that are typical of more traditional 2-line QTL studies. In addition, in the *D. melanogaster* system, a specialized piece of equipment, the “heat box”, originally developed by Wustmann et al. ^53^, allows for a high-throughput assay of place learning and memory^33, 54^. Place learning is a type of operant learning that occurs when a fly learns to associate a specific place with a consequence. In the heat box, up to 16 individuals can be assayed concurrently for place learning and memory in less than 10 minutes, making it feasible to assay the large number of individuals necessary for QTL mapping for these key phenotypes.

In this study, we use the DSPR to genetically dissect place learning and memory. After assaying 39,392 individual flies from 741 RILs from the DSPR, we mapped 16 QTL, with 5 shared QTL between learning and memory. In addition, we performed RNA-Seq to measure differential gene expression of high versus low performing cohorts of RILs and used these data to narrow the set of candidate genes within our QTL intervals. The loci we identify have not been previously associated with place learning or memory, representing a step forward in understanding the genetic basis of these traits.

## Methods

### Mapping population

We used a multi-parental population of *D. melanogaster,* the DSPR ^44, 51, 52^ (http://FlyRILs.org). The DSPR consists of two sets of approximately 800 RILs (genotype), each created via an 8-way advanced intercross design. To create these RILs, eight inbred founder lines were crossed and allowed to randomly mate for 50 generations followed by 25 generations of inbreeding. This creates a set of recombinant inbred lines whose genomes are mosaics of the eight original founder lines. Each founder line consists of a single inbred lines from a different location across the world, including North America, South America, Africa, Europe, and Asia ^51^, capturing some of the worldwide genetic variation in *Drosophila melanogaster*, though the amount of variation that can be captured by eight genotypes is necessarily limited. The founder lines have been fully re-sequenced, and the RILs have been genotyped at over 10,000 SNPs. King et al. ^44^ developed a hidden Markov model that infers the likely founder ancestry at all genomic locations for all RILs, providing full genome information for all RILs. Essentially, the hidden Markov model uses the genotypes at the ∼10,000 SNPs and the genome sequences of the founder lines to assign the most likely ancestry for each genomic segment in each RIL. King et al. ^44, 51^ give the complete details of the formation of the DSPR and associated resources. In this experiment, we used nearly all of the lines in one of the two sets of RILs, the population A RILs, which consisted of 741 RILs.

### Fly husbandry

All stock flies were raised on the standard cornmeal-yeast diet, and kept at 18° C, with 60% relative humidity, in wide vials (Fisher Scientific, Cat. No.: AS-513) until they were ready to be used in an experiment. Two weeks before the behavioral assays, about 6 males and 10 female flies were flipped onto a new flask (Fisher Scientific, Cat. No.:AS-355) of food and allowed to mate at 25°C and lay eggs. Seven days post-oviposition, all the adults were removed from the flask to ensure that only the F1 generation remained in the flask. Fourteen days post-oviposition, adult flies were anesthetized on ice, and we collected 60-80 female flies for the behavioral assay. These female flies were placed onto two separate new food vials in groups of 30-40 individuals per vial. The flies were then allowed to recover for at least 24 hrs before the assay. Flies included in the assay ranged in age from 15 to 22 days post-oviposition and, because they were reared with males, they were all presumably mated. We chose to phenotype only females in this experiment due to the existence of a large eQTL dataset in the DSPR using female heads. While ideally we would also phenotype males, this would have doubled the size of the experiment and was not feasible at the time. For each RIL we phenotyped between 31 and 134 individuals, with an average of 53 (Figure S1). It was not practical to perform this experiment with a fully blocked design, due to the number of RILs we assayed. In any given week, a total of 10-15 RILs were set-up and phenotyped. To assay such a large number of individuals per RIL, we also needed to perform assays on different days and at different times. All RILs were assayed across different days and different times to ensure there were not systematic differences between lines in the time of day and day of the week they were assayed. Across our entire dataset, the time of assay explained little variation in learning or memory, considering the month (Learning: R^2^ = 0.01; Memory R^2^ = 0.002), day of the week (Learning: R^2^ = 0.003; Memory R^2^ = 0.0006), or time of day (Learning: R^2^ = 0.005; Memory R^2^ = 0.003).

### Phenotyping

Through a behavioral assay known as place learning, we are able to train flies with a highly sensitive apparatus, the “heat box” ^53, 55, 56^. Single flies are placed in individual chambers that are lined top and bottom with Peltier elements, which allow for fast temperature changes within a chamber. The cooling (24°C), and warming (41°C) of the chambers – which takes about 4 seconds to change from one temperature to the other – are entirely controlled by the position of the fly. If the fly is positioned on the cool-associated side (24°C), then the entire chamber reflects that temperature, the same is true for the hot-associated side (41°C). There is an infrared light within each chamber that tracks the position of the flies during the assay. Because 41°C is a highly aversive stimulus, a typical fly will learn to remain on the cool-associated side of the chamber. In addition, most flies will remember to remain on the cool-associated side of the chamber once the punishment is removed (i.e., the chamber no longer heats when the fly enters the hot-associated side).

Using the above paradigm, we assayed learning and memory in a high throughput way. When all chambers are functioning, a total of 16 flies can be tested within the heat box concurrently, with both learning and memory assays performed consecutively on a single individual fly. In another type of assay of place learning in flies, it has been shown that the performance of flies depends on whether they are assayed alone or in groups ^57, 58^. While the heat box assays up to 16 flies at once, they are each assayed in an individual chamber. These chambers are entirely dark and are each housed in a separate box within the larger heat box, making it unlikely flies receive sensory information from one another during the assay. We define learning as the increasing avoidance of the hot-associated side, and memory as the persistent avoidance of the hot-associated side, even if the aversive stimulus no longer exists ^1–5^. To quantify learning and memory, flies are given a total of 9.5 mins within the chamber: 30 secs for pre-testing, 6 mins for learning, and 3 mins memory testing. During these testing periods, performance indices (PIs) were calculated with a specialized software (Heat Calc 2.14 ^54^), which reflect the relative position preference of an individual fly. We calculated PIs for both phenotypes, using the amount of time an individual spends on one side of the chamber divided by the total amount of time of the assay (Figure 1):

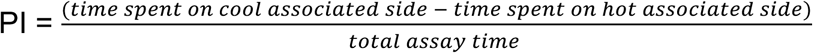

**Figure 1:**
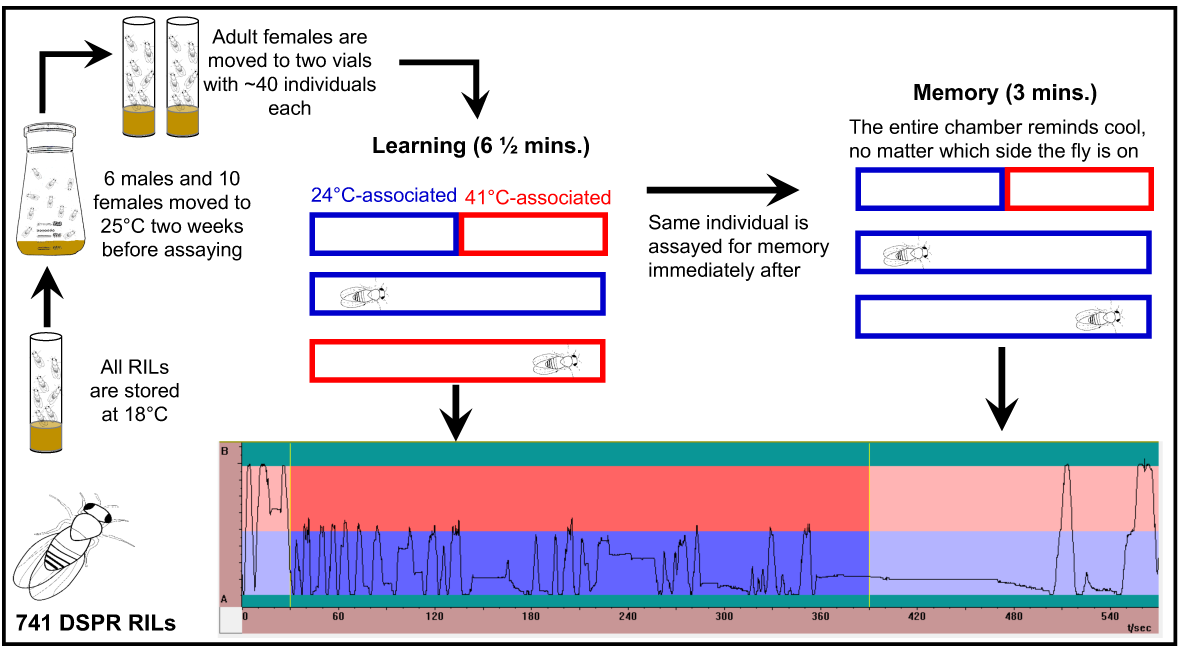
Schematic of phenotyping process of 741 RILs (39,392 individuals). RILs are stored at 18°C, until 1 day of “set up”, then 6 males and 10 females are placed into a flask and allowed to mate. Fourteen days post-oviposition adult F1 flies were collected into two separate vials. These flies were then phenotyped within the heat box. The rectangles depict a chamber within the heat box, blue (cool) and red (hot) represent 24°C and 41°C respectively, which are the temperatures that are associated with one half of the chamber. If an individual is on the cool associated side of the chamber, then the whole chamber reflects that temperature, and the same is true for the hot associated side. After 30 s pre-testing, learning is phenotyped for 6 mins., and immediately after the same individual is phenotyped for memory for 3 mins. Traces are the activity within a chamber of an individual fly. This individual has a higher learning score, because it spent less time of the hot associated side, whereas the memory score is medium.

The maximum PI is 1, which corresponds to a fly that has perfect avoidance of the hot-associated side. A PI of zero indicates preference for neither side, and a PI of −1 indicates a fly that remains on the hot-associated side for the entire assay.

To ensure our measurements of learning and memory do not actually reflect variability in activity or thermal tolerance, we also assayed activity and thermal tolerance for each individual. Prior to the start of the learning and memory assays, we track each fly’s activity for 30 seconds to estimate individual activity. Additionally, after the learning and memory assays, we assayed the time to incapacitation when the chamber is held at a constant 41°C as a measure of thermal tolerance. For each of these phenotypes, we fit a linear regression to estimate the amount of variation in learning or memory that is accounted for by activity or thermal tolerance. Thermal tolerance explains little variation in either learning (*R^2^* = 0.002) or memory (*R^2^* = 0.005). Activity explains more, but still a modest amount of variation, in learning (*R^2^* = 0.07) and memory (*R^2^* = 0.02). Thus, it is unlikely our measures of learning and memory are driven by variability in activity or thermal tolerance. In addition, the trend that does exist points to more active individuals having lower learning and memory scores (Figure S2 & S3), which is contrary to what one would expect if our results were the product of variability in the general health of the lines because you would expect healthier lines to both learn better and be more active.

Several quality control filters allowed us to remove any individual assays with problems and ensure the validity of all measurements. These analyses and all following were performed in R ^59^. First, the Heat Calc software eliminates any individuals that did not cross the midline of the chamber at least once, to ensure all individuals experienced the aversive stimulus. Second, the temperature of each chamber is tracked over the course of the assay, and we used these data to identify problematic chambers (i.e., any chamber that was not warming or cooling to the correct temperature). Most often, these were chambers that were not reaching the target temperature of 41°C. We flagged any chamber as problematic if the average difference between the actual temperature within the chamber and the target temperature exceeded 5°C during the assay. Of a total of 48,943 assays, we eliminated 8,688 via this criteria, the vast majority of which represented two chambers that at the time were unknowingly failing to reach the appropriate target temperature. We identified the issue with these chambers via this quality control analysis. Finally, we eliminated any individual that showed a long period of inactivity at the end of the assay to remove any individuals that were potentially immobilized for any reason, which would skew their learning or memory PI. This criteria occurred for only 863 assays, and it was not specific to a certain set of RILs, chambers, or assay days.

### Heritabilities and QTL mapping

We estimated the broad sense heritability of learning and memory and the genetic correlation between these phenotypes by estimating the genetic and phenotypic variance and covariance components from a linear mixed model using the `lme` and `VarCorr` functions in the `nlme` package ^60^, followed by a jackknife, which allowed us to obtain standard errors of our estimate ^61, 62^. Prior to fitting this model, both learning and memory were quantile normalized to ensure normality. This transformation was particularly important for the individual-level data because with the PIs bounded by −1 and 1, the data are highly non-normal. The jackknife removes each observation once, the model is then fit and the quantitative genetic parameter (e.g., heritability or the genetic correlation) is estimated, and a pseudovalue is calculated. Thus, for our dataset of 741 RILs, each RIL is deleted, one at a time, to produce 741 pseudovalues. Because the `lme` function uses restricted maximum likelihood, it is possible for the model to not converge. This occurred for 28 cases for the heritability of learning, 50 cases for the heritability of memory, and 77 cases for the genetic correlation, thus, these pseudovalues were not included in the calculation of our estimates. Our estimate did not differ substantially from the estimate obtained from fitting the mixed model alone without using the jackknife. We also performed a model comparison using a likelihood ratio test to determine the significance of RIL (i.e., genotype). We compared the model fitting only the intercept to the model including RIL, fitting both via maximum likelihood such that model comparisons would be valid.

For each of the 741 RILs we measured, we took the average PI for learning and memory from each RIL, and applied Haley-Knott regression ^43^. This statistical approach regresses each phenotype on the eight founder haplotype probabilities ^43, 44^ by fitting the following model:

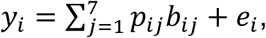

where *y_i_* is the phenotype (i.e., learning or memory) of the ith RIL, *p_ij_* is the probability that the *i*th RIL has the *j*th haplotype at the locus, *b_ij_* is the vector of effects for the *j*th haplotype, and *e_i_* is the vector of residuals. We fit this model at ∼10,000 regularly spaced positions across the genome. Prior to performing this genome scan we transformed the learning phenotypic data, using a power transformation, raising our average learning values to the third power, which improved the normality of the data to meet the assumptions of our statistical test (Figure S4). The memory data was approximately normally distributed and did not require a transformation. We also performed QTL mapping on the residuals of 1) a model correcting for individual activity and 2) a model correcting for individual thermal tolerance and show that our results are largely similar to our uncorrected results in both cases (Figure S5 & S6). The resulting p-values are highly correlated between the raw and corrected values (Activity: Learning: r = 0.84; Memory r = 0.90; Thermal tolerance: Learning: r = 0.97; Memory r = 0.96). Given these results, we favor the raw learning and memory measurements, because the analysis and interpretation is more straightforward. To identify statistically significant QTL, we performed a 1000 permutations ^63^ of the dataset and performed a genome scan on each permuted dataset as above. The same individuals were assayed for learning, memory, and thermal tolerance. We do not report the results of thermal tolerance here; however, because all three phenotypes are part of the same experiment, we permuted these three phenotypes together and performed genome scans at once. We then determined the number of false positive QTL at different significance thresholds. We identified individual, distinct QTL by removing any QTL that were within 2 cM of a more significant QTL. We calculated the empirical false discovery rate (FDR, expected false positives/total positives) for each threshold and determined the threshold corresponding to a 5% FDR. We also calculated the threshold corresponding to a 5% family-wise error rate (FWER, the probability of one or more false positives experiment wide) by determining the lowest p-value for each set of genome scans from each permutation and calculated the 95% quantile of the resulting set of p-values ^63^. For each peak, we calculated the estimated effects of each haplotype at each QTL, the percent variance explained by the QTL (PVE), and the Bayesian Credible interval (BCI), following the methods described previously for the DSPR ^44^. We considered QTL as separate QTL if their BCIs did not overlap. One memory QTL, Q15 is near a wider learning QTL, Q14, and thus it is possible Q15 is a shared, pleiotropic QTL. However, within learning, the peak at Q14 and a smaller peak near Q15 cannot be distinguished as their BCIs overlap. Additionally, while Q14 and Q15 are near one another, their BCIs do not overlap. Thus, we decided to be conservative with respect to concluding Q15 was a shared QTL and consider it a QTL for memory only.

### Rna-sequencing and Analysis

We selected 14 RILs from the high (n = 7) and low (n = 7) 5% for learning and memory of the first 140 RILs we phenotyped. Since the maximal and minimal PIs are bounded by 1 and −1, we felt confident the RILs in the top and bottom cohorts would still have higher or lower than average PIs after assaying all RILs. Obviously, the ranking of these RILs changed once we assayed an additional 601 RILs, however, they do still fall within the high and low cohorts. We collected 25 female heads from each of the 7 RILs in the high and low cohorts. Five heads from each RIL within a cohort were pooled into 5 biological replicates, totaling 35 heads per tube. To optimally identify consistent differences in expression between high and low cohorts, we chose to pool RILs together to average over biological differences and reduce the number of samples for RNA-Seq. We also included 5 heads per RIL to account for individual-level variability.

To prepare the samples, first the 7 selected RILs underwent the same set-up protocol as done in the phenotypic assay (see phenotyping section above for details). Briefly, female flies (fourteen days post-oviposition) were selected by anesthetizing them on ice between 0800 and 1000 and allowed to recover for 24 hrs at 25°C. Immediately after, the flies were flash frozen in liquid nitrogen. Second, the samples were vortexed for no more than 10 seconds to remove heads from bodies and stored at −80°C. Third, the samples were freeze dried overnight to prevent degradation using a lyophilizer (Labconco, Cat No.: 77550-00). Both the high and low cohorts were processed in the same batch for each phenotype.

The samples were then shipped to RaPiD Genomics, LLC (fee for service) where the total RNA was extracted with Dynabeads mRNA direct kit from life technologies, mRNA was then fragmented and converted into double stranded cDNA, followed by the standard proprietary library prep for one lane of Illumina HiSeq 3000 instrument to generate paired-end (PE) reads. The first 16 samples were PE 150-bp, while the remaining samples were PE 100-bp.

Before aligning the reads, we trimmed the samples using the software *cutadapt* ^64^, then ran a quality control test using *fastqc* ^65^. The fastqc analysis did not reveal any major problems with the reads. We specifically assessed the summaries for per base sequence qualities and the per sequence quality scores. We aligned reads to the *D. melanogaster* reference transcriptome (Ensembl Release Version: 84) using HISAT2 ^66^ and assembled and quantified transcripts using StringTie ^67, 68^. We did not allow the assembly of novel transcripts not present in the *D. melanogaster* reference. We used the “prepDE” python script available from the StringTie manual to calculate the read counts for each gene. To assess differentially expressed genes between our high and low cohorts, we used the DESeq2 package ^69^. First, we filtered all genes that had zero counts in all samples. We then performed surrogate variables analysis (SVA) on normalized counts using the sva package ^70, 71^ to estimate for unknown batch effects while accounting for treatment. The SVA identified one surrogate variable, which we included as a covariate when testing for differentially expressed genes. To identify significantly differentially expressed genes we used the DESeq2 package ^69^ to test for overall treatment effects. We then performed contrasts between the high and low cohorts of learning and the high and low cohorts of memory to identify significantly differentially expressed genes. To visualize and rank the genes we used the function `lfcShrink`, which performs shrinkage on log_2_(Fold Changes), which have been shown to produce better estimates. All log_2_(Fold Changes) reported here are the shrinkage estimated values using the `“normal”` estimator.

To identify possible candidate genes within our QTL intervals, we first determined which of the genes within each interval of interest were significantly differentially expressed between high and low performing cohorts. For QTL mapped to a single phenotype, this interval was the BCI. For QTL mapped to both phenotypes, we considered only the overlap region between the BCI’s for both learning and memory. In addition, we integrated a previous genome-wide eQTL dataset using the DSPR. King et al. ^72^ performed eQTL mapping on ∼600 RIL crosses, crossing the population A and population B RILs and measuring expression of nearly all genes in the *D. melanogaster* genome using microarrays. We used this dataset to identify which genes show evidence for a *cis* eQTL in the DSPR. Finally, we identified differentially expressed genes with previous evidence implicating a potential role in learning and memory by manually examining the annotation for these genes in FlyBase ^73^ and noting annotations such as neurological process, behavior, or neurotransmitter, which might implicate the gene could be involved in the process of learning or memory.

### Data Availability

Raw learning and memory phenotypic data are available from zenodo (https://zenodo.org/): http://doi.org/10.5281/zenodo.2595557. RNA-Seq data is available from the NCBI Short Read Archive ^74^ under SRA accession number: PRJNA527143 (https://www.ncbi.nlm.nih.gov/sra/PRJNA527143). Raw re-sequencing data of the founder lines is deposited in the NCBI SRA under accession number SRA051316, and the RIL RAD genotyping data are available under accession number SRA051306. Founder genotype assignments from the hidden Markov model are available as two data packages in R (http://FlyRILs.org/) and are available from the Dryad Digital Repository (http://dx.doi.org/10.5061/dryad.r5v40). See King et al. ^44, 51^ for details of the DSPR datasets. All data associated with the DSPR eQTL dataset is also available publicly (see ^72^ for experimental details) in NCBI’s Gene Expression Omnibus ^75^ and are accessible through GEO Series accession number GSE52076. All of the above DSPR data are also available centrally at http://FlyRILs.org. The complete code used to perform all analyses is available at GitHub (https://github.com/EGKingLab/LearnMem_DSPR).

## Results

### Phenotypic Patterns

We measured place learning and memory in 741 RILs from the DSPR (Figure 1) and show these lines vary widely in both these traits (Figure 2). For each line, we measured a minimum of 31 females, with an average of 53 per line (Figure S1). The lines range in average learning performance index (PI) from 0.215 to 0.933 from the lowest performing RIL to the highest, while average memory PIs range from −0.016 to 0.874 (Figure 2). The estimated broad-sense heritabilities are moderate (Learning: H^2^ = 0.20, 95% CI = 0.18–0.22; Memory: H^2^ = 0.10, 95% CI = 0.09–0.12), and the effect of RIL (i.e., genotype) is highly significant (Learning 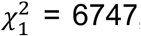, p < 0.0001; Memory 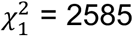, p < 0.0001), demonstrating a genetic basis for these traits. Learning and memory are genetically correlated (*r_g_* = 0.53, 95% CI = 0.47–0.59; Figure 2), indicating that some of the same loci influence both learning and memory. This relationship would be expected given memory formation cannot occur without learning, though high learning performance does not necessarily guarantee high memory performance. Indeed, while most RILs with high learning PIs also have high memory PIs, some RILs show high learning but poor memory, indicating some independence between these processes as well.

**Figure 2:**
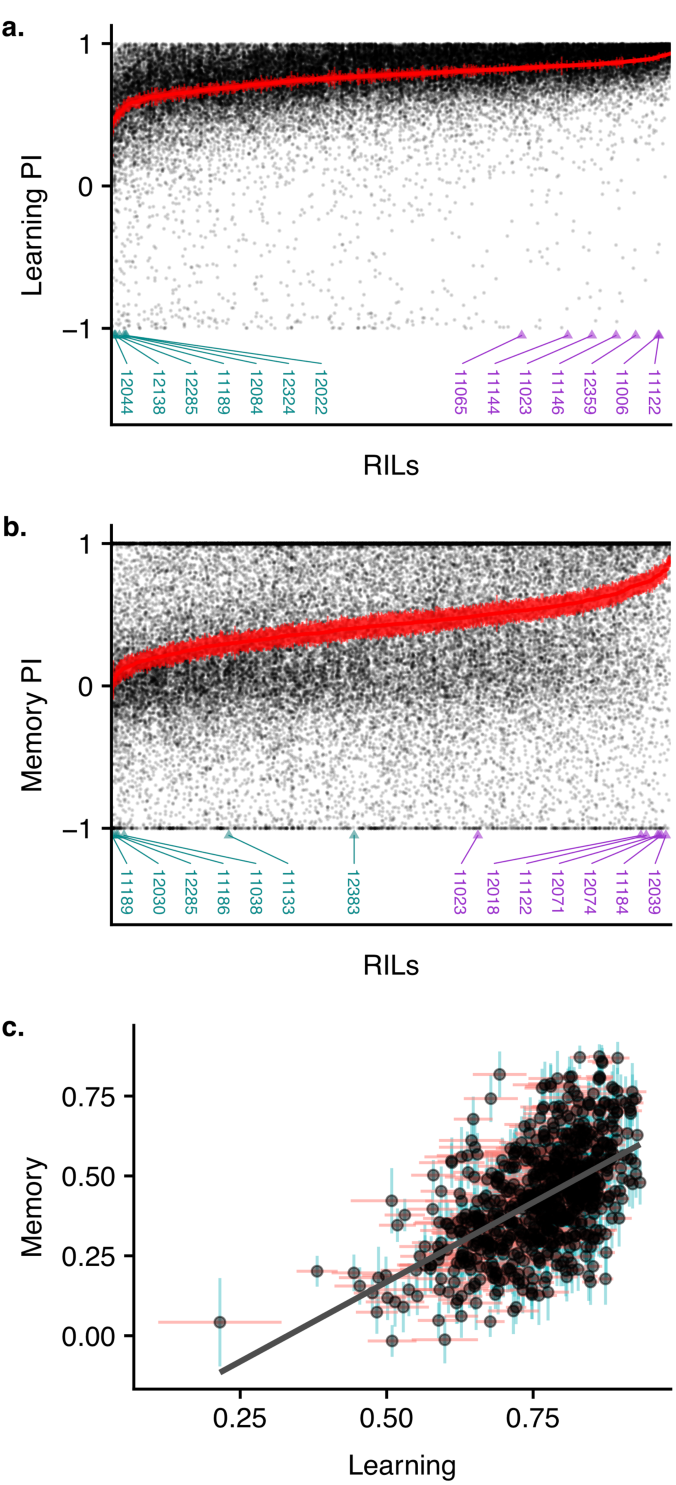
Phenotypic patterns in place learning and place memory among the DSPR RILs. a) Performance index (PI) scores for learning for each RIL in the DSPR. The RILs are sorted from lowest average learning to highest average learning. The PIs for each individual measured for a given RIL are displayed (black points) and the mean value ±1 SE for each RIL is plotted in red. RILs selected for RNA-Seq in the low performing cohort are labeled with cyan triangles, and the high performing cohort is labeled with purple triangles. b) PI scores for memory for each RIL in the DSPR plotted in the same way as learning in (a). c) The relationship between learning and memory. The mean values for learning and memory for each RIL are plotted with black points. Horizontal red bars show the learning mean ±1 SE and the vertical blue bar shows the memory mean ±1 SE.

### Genomic Scans (QTL)

To identify loci affecting place learning and memory, we performed a genome scan following established methods for mapping QTL in the DSPR ^44, 51^. To establish both a 5% family-wise significance threshold (FWER) and a 5% false-discovery rate threshold (FDR), we performed permutations (see Methods for details). We identified 16 QTL (Q1 - Q16) at a 5% FDR: 9 are unique to the learning phenotype, 2 are unique to the memory phenotype, and 5 were mapped for both phenotypes (Figure 3; Table 1). Of these, 10 QTL are also significant at the FWER threshold (Table 1). For each QTL, we obtained a 95% Bayesian Credible Interval (BCI) ^43, 76^ to define the region in which the causative variants are expected. With the exception of Q12 for learning, which spans the centromere, these intervals are narrow, averaging 643 kb in physical distance and 2 cM in genetic distance. All of the QTL we identified are of moderate effect, with the percentage of phenotypic variance explained by a locus averaging 4% and ranging from 3% to 7% (Table 1). In the DSPR, we are able to assign the likely haplotype identity to every genomic segment in every RIL ^44^. Therefore, for our mapped QTL, we can estimate the effect of harboring each of the 8 founder haplotypes at the QTL location on our phenotypes of interest. We show these estimates, along with the haplotype assignments for each individual RIL measurement for all shared QTL in Figure 4. As has been found previously in the DSPR ^72, 77^ and in other mapping panels ^78, 79^, these effect estimates show evidence for multiple causative variants resulting in an allelic series, not a single causative biallelic SNP that would show haplotype effects in defined “high” and “low” groups. The similar patterns in our effect estimates for learning and memory for a single shared QTL also support the hypothesis that these are pleiotropic loci, rather than separate linked QTL, which would be expected to show contrasting patterns in the two phenotypes.

| ID | Chromosome | Traits(s) | Position (Mbp) | Lower (Mbp) | Upper (Mbp) | -log<sub>10</sub>(P-value) | PVE (%) | Hypothesized candidate genes<sup>b</sup> |
| --- | --- | --- | --- | --- | --- | --- | --- | --- |
| Q1 | X | Learning | 7.02 | 5.79 | 7.69 | 3.25 | 3.43 | - |
| Q2 | X | Learning | 15.75 | 15.24 | 16.74 | 3.20 | 3.40 | - |
| Q3 | X | Learning | 18.81 | 18.72 | 19.06 | 4.71<sup>a</sup> | 4.48 | CG7058 |
| Q4 | 2L | Shared | L: 3.19 | 3.06 | 3.22 | 5.58<sup>a</sup> | 5.07 | Mad |
|  |  |  | M: 3.16 | 3.04 | 3.25 | 3.66 | 3.73 |  |
| Q5 | 2L | Shared | L: 7.59 | 6.75 | 7.77 | 4.19 | 4.12 | - |
|  |  |  | M: 7.48 | 7.47 | 7.61 | 5.25<sup>a</sup> | 4.85 |  |
| Q6 | 2R | Memory | 10.16 | 9.70 | 10.35 | 6.63<sup>a</sup> | 5.78 | - |
| Q7 | 2R | Learning | 18.68 | 18.51 | 19.35 | 4.28 | 4.18 | - |
| Q8 | 2R | Learning | 22.34 | 22.28 | 22.46 | 5.12<sup>a</sup> | 4.76 | Liprin-γ; wdp |
| Q9 | 3L | Shared | L: 1.68 | 1.20 | 2.71 | 4.13 | 4.07 | mwh |
|  |  |  | M: 1.30 | 0.82 | 1.36 | 3.56 | 3.67 |  |
| Q10 | 3L | Learning | 6.92 | 6.59 | 7.13 | 5.31<sup>a</sup> | 4.89 | Mp |
| Q11 | 3L | Shared | L: 8.57 | 8.50 | 8.58 | 5.89<sup>a</sup> | 5.28 | ergic53 |
|  |  |  | M: 8.61 | 8.31 | 8.90 | 3.52 | 3.63 |  |
| Q12 | 3L | Shared | L: 15.64 | 3L:15.41 | 3R:4.52 | 3.44 | 3.58 | - |
|  |  |  | M: 15.66 | 15.64 | 15.67 | 4.71<sup>a</sup> | 4.48 |  |
| Q13 | 3R | Learning | 19.60 | 19.46 | 21.32 | 4.17 | 4.10 | - |
| Q14 | 3R | Learning | 22.45 | 22.43 | 22.68 | 7.78<sup>a</sup> | 6.53 | Octβ1R |
| Q15 | 3R | Memory | 23.06 | 23.02 | 23.15 | 4.37 | 4.24 | - |
| Q16 | 3R | Learning | 27.55 | 27.48 | 27.86 | 6.89<sup>a</sup> | 5.95 | CG34353 |

<sup>a</sup>These QTL significant at the more stringent 5% FWER. All QTL in the table are significant at a 5% FDR
<sup>b</sup>Hypothesized candidate genes under the QTL peak. A gene is only listed if there are multiple sources of evidence for the gene. See text for details.

**Figure 3:**
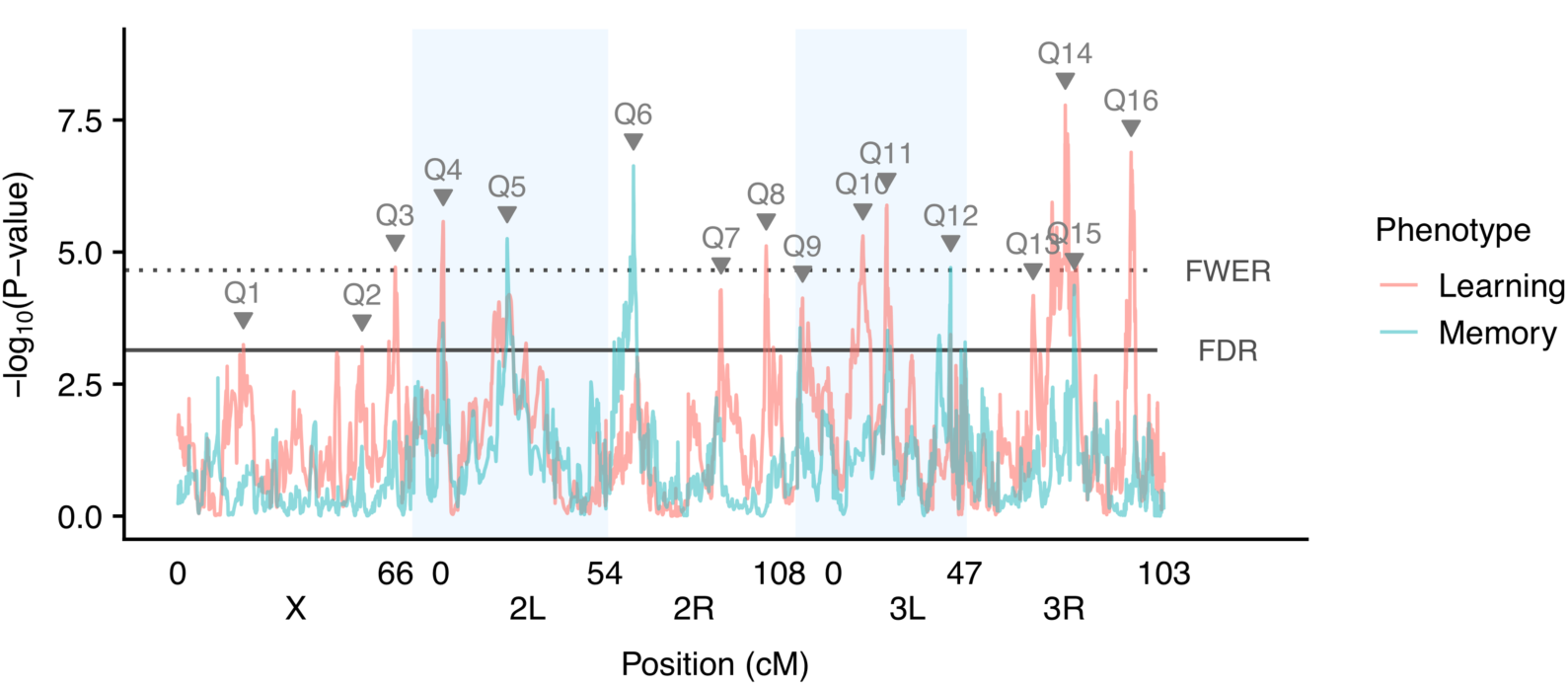
Genome scan of learning and memory phenotypes. The −log_10_(P value) versus position for both learning and memory. Taller peaks represent QTL with stronger statistical support. Different colors denote the different phenotypes of learning and memory. Dotted horizontal line denotes the 5% family-wise error rate threshold and the solid horizontal line denotes the 5% false discovery rate. QTL peaks reaching either threshold are labeled with the QTL id. Shaded blue boxes denote chromosome arms.

**Figure 4:**
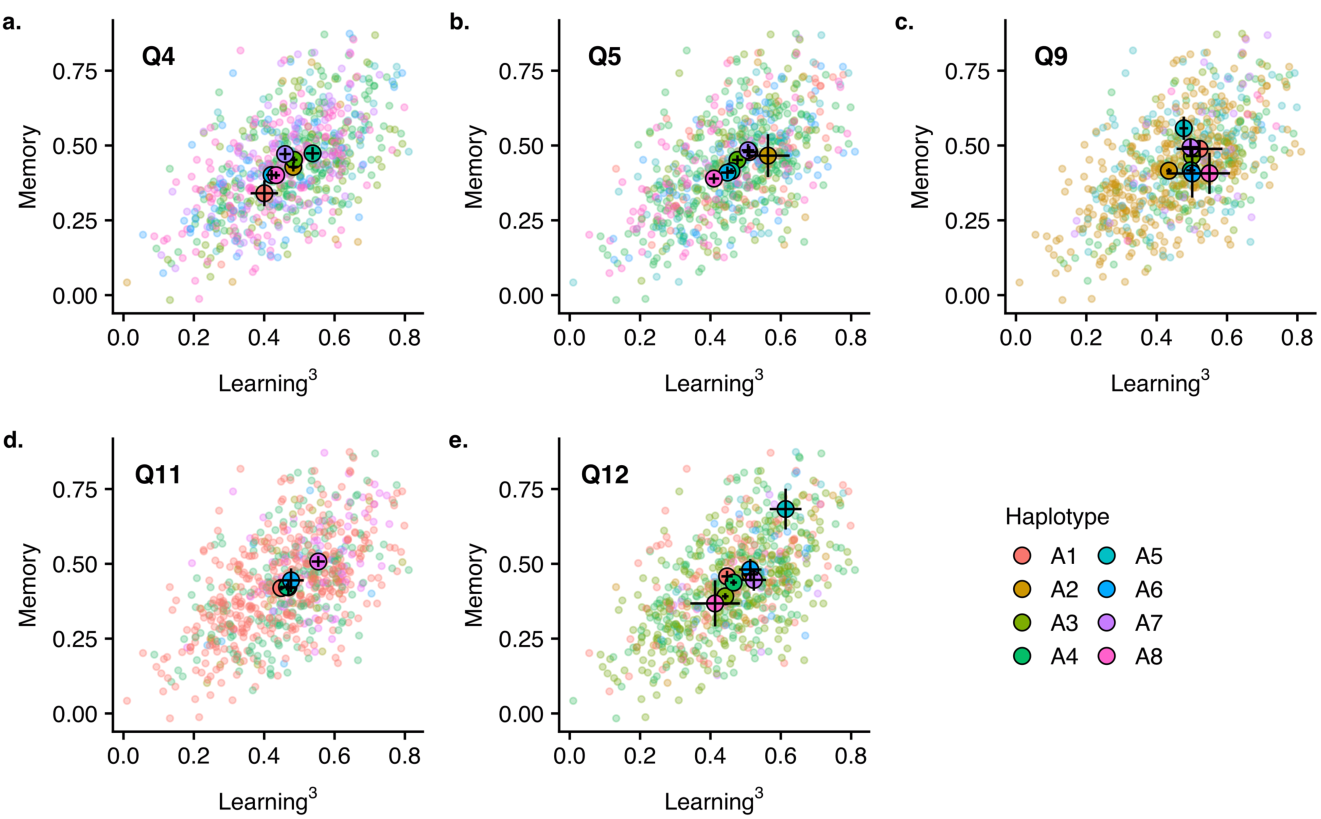
Haplotype means at each shared QTL. The average learning (power transformed) and memory PI are plotted for each RIL. Each RIL is colored by its most likely haplotype identity at the location of the QTL. The large black outlined points are the estimated average effect of each haplotype and the horizontal and vertical bars show ±1 SE of this estimated effect for learning and memory respectively. The individual RILs are plotted with semi-transparent points.

### Gene Expression

We performed RNA-Seq on pooled sets of female heads to identify differentially expressed genes in cohorts of high or low performing RILs for place learning and memory. For each, we selected 7 high performing and 7 low performing RILs and pooled a total of 35 female heads in a single sample, with 5 heads from each RIL contributing to each sample (Figure 2). We then performed RNA-Seq on 5 biological replicates of these pooled samples (see Methods for details). This design allowed us to account for individual- and RIL-level variability while keeping the number of RNA-Seq samples reasonable. The resulting dataset provides a genome-wide expression profile of high and low performers for place learning and memory (Figure 5). For place learning, 2076 genes were differentially expressed between the high performing and low performing cohorts. Of these, 947 genes were upregulated and 1129 were downregulated in the high learning cohort relative to the low learning cohort. For place memory, 590 genes were differentially expressed between the high and low performing cohorts, with 347 upregulated genes and 243 downregulated genes in the high memory cohort relative to the low memory cohort. There are 220 genes that are significantly differentially expressed in both the learning dataset and the memory dataset, the majority of which (n = 169) trend in the same direction for both phenotypes. There are several genes, including *dunce, amnesiac, white, radish, rutabaga, arouser, and tribbles*, have been previously implicated as playing a role specifically in place learning and memory ^80^, which are labeled on Figure 5. Of these, only *radish* is significantly differentially expressed between high and low learning cohorts, with higher expression in the low learning cohort (log_2_FC: −0.4, p_adj_ = 2.4 x 10^-5^). Comparing memory cohorts, only *dunce* and *white* were significantly differentially expressed, and both were more highly expressed in the high memory cohort (*dunce:* log_2_FC: 0.38, p_adj_ = 0.005; *white*: log_2_FC= 0.6, p_adj_ = 1.6 x 10^-5^). We note that *white* is a commonly used marker and is often assumed to not affect the phenotype being interrogated in these studies, an assumption that our results and others challenge. Here, we show it varies between high and low memory cohorts and note it has been previously found to influence place learning and memory^81^, as well as courtship behavior in males^82^.

**Figure 5:**
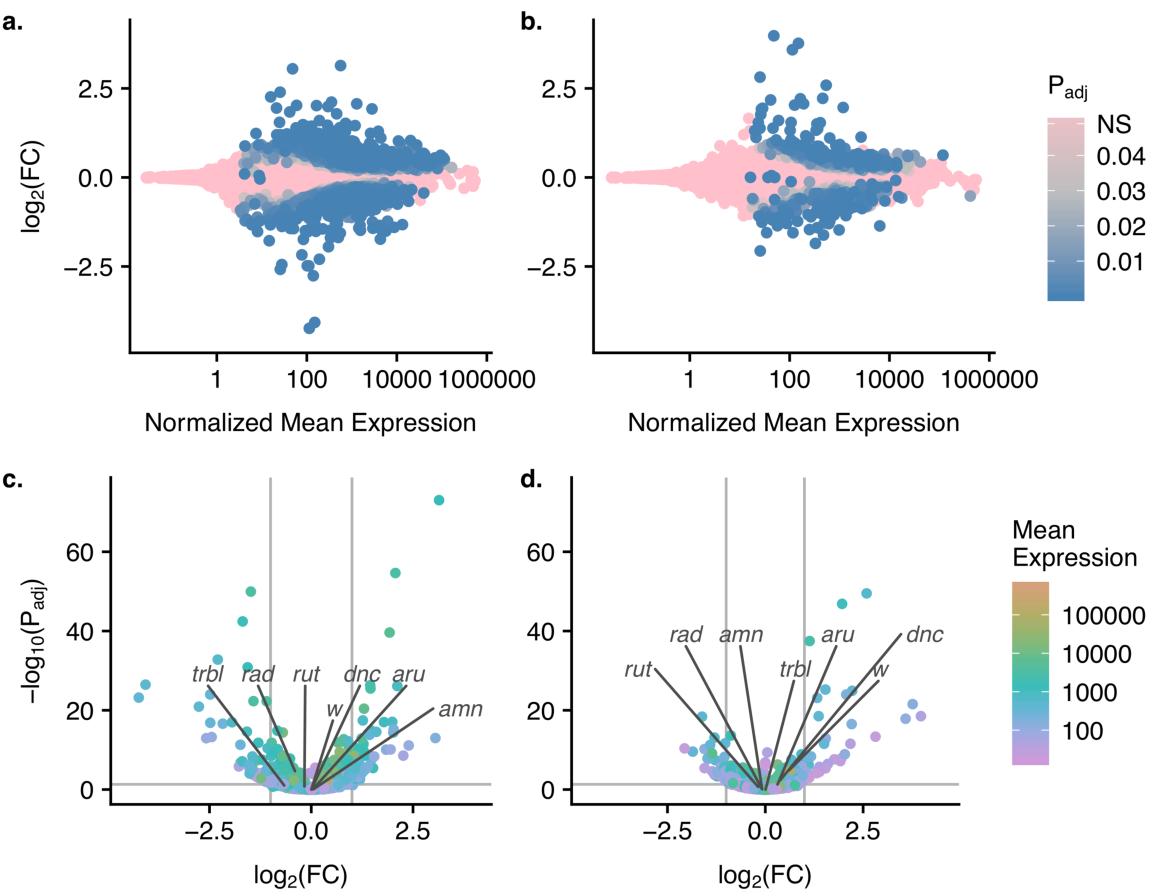
RNA-Seq expression levels between high and low cohorts of learning and memory in the DSPR. a, b) MA plot of learning cohorts (a) and memory cohorts (b) showing the log_2_FC versus the normalized average expression for each gene. Colors indicate the adjusted p-value for a given gene. c, d) Volcano plots of the comparison between high and low performing cohorts for learning (c) and memory (d). Log_2_FC are shrinkage adjusted and represent the high performing cohort relative to the low performing cohort. Colors indicate the overall average expression level. The points corresponding to a set of genes that have been previously associated with place learning and memory are labeled.

### Identification of candidate genes

While our QTL credible intervals are fairly narrow, they do not provide the single gene resolution that would be necessary to immediately identify a potential causative gene. We used our RNA-Seq data to identify potential regulatory candidate genes within our QTL BCI. For QTL shared between learning and memory, the causal gene should lie within the overlapping region of the learning and memory BCIs, assuming these QTL truly represent a shared causal gene. This assumption allowed us to further narrow the search intervals for these shared QTL. We then determined which significantly differentially expressed genes fall within the interval of interest for the 16 identified QTL. We used two additional sources to inform the identification of potential candidate genes. First, the DSPR is a widely used resource, providing datasets that have the potential to integrate with this study. A previous study by King et al. ^72^ performed genome-wide eQTL mapping of female heads for ∼600 RIL crosses. These RILs were crosses between the population A RILs and the population B RILs, while our assays only used the population A RILs. Nevertheless, this dataset allows us to identify which genes within an interval have been previously identified as having a significant *cis* eQTL in the DSPR. Second, we examined the FlyBase (FB2018_06) controlled vocabulary terms associated with the genes in our QTL intervals to identify any genes with a potential role in learning or memory ^73^. Each QTL BCI is shown in Figure 6, with the significantly differentially expressed genes within the interval labeled, except for Q12, which spans the centromere and thus has an interval too wide to display effectively. Some QTL are fairly wide, such as Q1, Q2, Q6, Q7, and Q13, and thus include a moderate number of differentially expressed genes with several as possible candidates. In one case, Q12, the interval of interest includes no significantly differentially expressed genes. For all of our other QTL, we were able to identify potential candidate genes.

**Figure 6:**
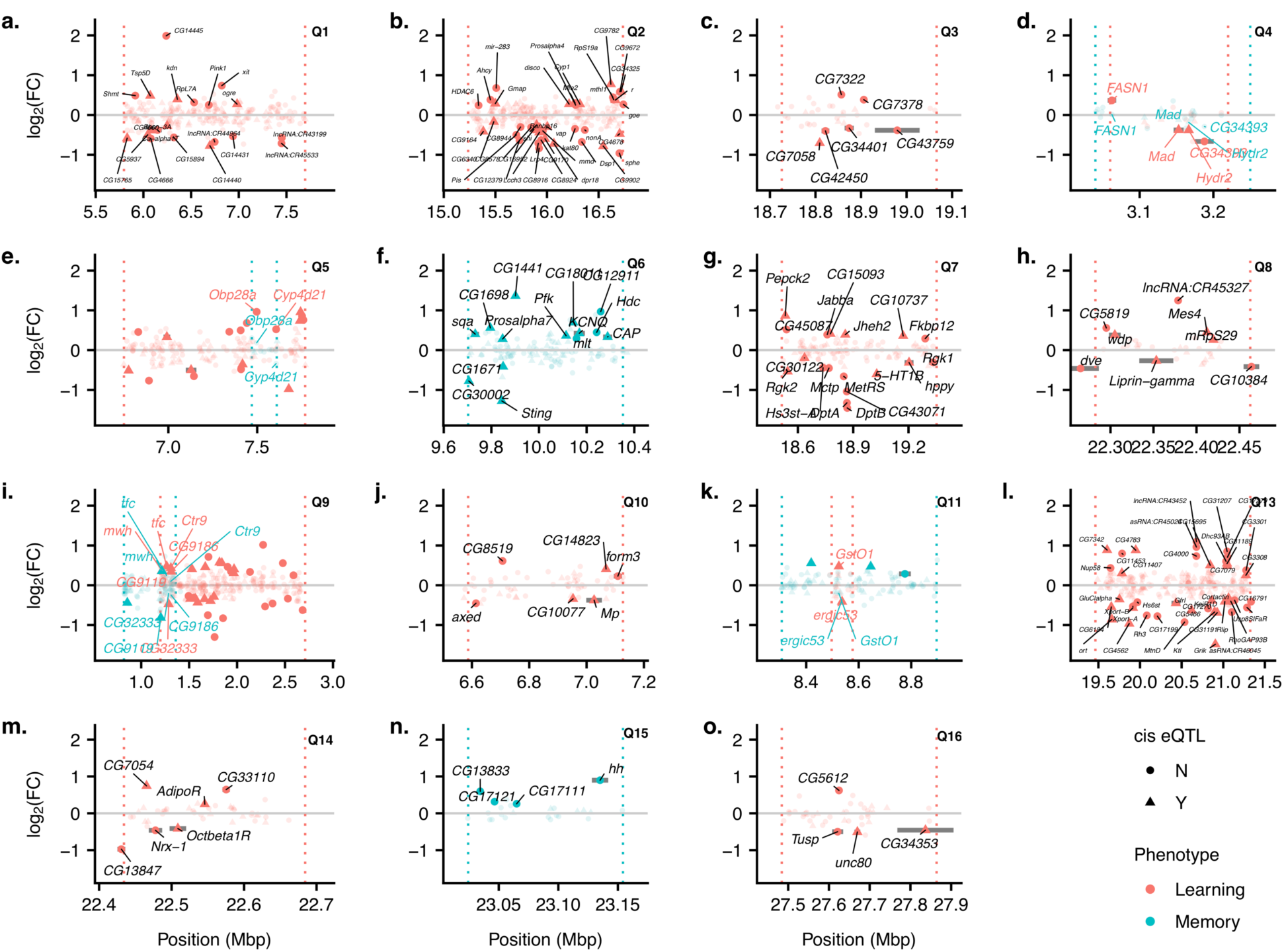
Individual QTL BCI regions showing possible candidate genes. Each panel from a-o shows one QTL BCI. When a peak is shared between learning and memory, both BCIs are shown. The dotted vertical lines denote the BCI limits on either side. Each point shows the location and the log_2_FC for a gene that occurs within the BCI. Significantly differentially expressed genes are plotted with solid points and non-significant genes are plotted with semi-transparent points. A given gene is labeled if it falls within the interval of interest (within the BCI for a single trait or within the region of overlap of BCIs if it is shared) and is significantly differentially expressed. Circles show genes that do not have evidence for a cis eQTL and triangles are genes that do have a significant cis eQTL. Colors denote the different phenotypes with red corresponding to learning and blue to memory.

Among the QTL mapped only for learning, Q3, Q8, Q10, Q14, and Q16 are narrow enough to suggest a small number of possible candidate genes. Only six genes that fall within the Q3 interval (Figure 6c) are differentially expressed, and just one has a *cis* eQTL in the King et al. ^72^ dataset: *CG7058*. The log_2_(Fold Change) (hereafter log_2_FC) shows *CG7058* is expressed at higher levels in the low learning cohort relative to the high learning cohort (log_2_FC= −0.7, p_adj_ = 9.3 x 10^-5^). Little is known about the function of this gene beyond it is known to be expressed in the nervous system ^83^. Of the eight genes that fall within the interval for Q8 (Figure 6h), four have a *cis* eQTL in the King et al. ^72^ dataset. Two of these genes have been shown previously to be involved in the nervous system. *Liprin-γ* is expressed at higher levels in the low learning cohort relative to the high learning cohort (log_2_FC= −0.27, p_adj_ = 0.016) and is one of three *Liprin* genes in the *D. melanogaster* genome that are involved in synapse formation. Astigarraga et al. ^84^ used null mutants of all three of the *Liprin* genes to show that *Liprin-α* and *Liprin-β* both promote synapse formation, and *Liprin-γ* acts antagonistically to both of these. In addition, *windpipe (wdp)* was more highly expressed in the high learning cohort relative to the low learning cohort (log_2_FC= 0.37, p_adj_ = 0.001). This gene has been implicated in synaptic target recognition in the neuromuscular junction via a screen in which *wdp* was overexpressed in the target tissue in larvae and the number of mistargeting events was quantified ^85^. In the Q10 (Figure 6j) interval, of the six differentially expressed genes, three have previous evidence for a *cis* eQTL ^72^. *Multiplexin (Mp)* is more highly expressed in the low learning cohort relative to the high learning cohort (log_2_FC= −0.38, p_adj_ = 0.029), encodes a Collagen XV/XVIII type protein, and has been shown to be critical for presynaptic homeostasis in the neuromuscular junction ^86^. Within the Q14 (Figure 6m) interval, three significantly expressed genes have previous evidence for a *cis* eQTL ^72^. One of these genes, *Octopamine β1 receptor (Octβ1R)*, is one of the octopamine receptors in flies and is more highly expressed in the low learning cohort relative to the high learning cohort (log_2_FC= −0.41, p_adj_ = 0.008). In insects, octopamine is a major neurotransmitter that has been shown to play a critical role in learning, particularly reward-based learning ^87–89^. A previous study by Sitaraman et al. ^90^ found no difference in place learning in flies with greatly reduced octopaminergic signaling, suggesting octopamine signaling is not necessary for place learning. However, Koon and Budnik ^91^ also showed that the *Octopamine β1 receptor (Octβ1R)* inhibits synaptic growth, acting in opposition to *Octopamine β2 receptor (Octβ2R*), which promotes synaptic growth. The *Octβ1R* inhibitory effect occurs via inhibition of the cAMP pathway, which is the same pathway affected by the well-known learning and memory mutants *dnc* and *rut* ^91^. Only four genes are significantly differentially expressed within the Q16 interval (Figure 6o), and two of these have a known *cis* eQTL ^72^. One of these, *CG34353*, has been classified with terms such as axon guidance and synapse organization via the PANTHER classification system ^92^, though these functions have not been confirmed in the *D. melanogaster* system. This gene is more highly expressed in the low learning cohort relative to the high learning cohort (log_2_FC= −0.46, p_adj_ = 0.01).

We mapped just two QTL specifically for our memory phenotype, and only Q15 (Figure 6n) was sufficiently narrow to identify possible candidate genes. There were just four significantly differentially expressed genes within this interval, none of which have a *cis* eQTL in the King et al. ^72^ dataset. All four of these genes are more highly expressed in the high memory cohort relative to the low memory cohort. The gene with the largest difference is the well-known *hedgehog* gene *(hh;* log_2_FC= 0.90, p_adj_ = 6.4 x 10^-5^), which has multiple, critical roles in development ^73^.

For the QTL mapped for both learning and memory, our ability to narrow the interval of interest to the region of BCI overlap between both traits leads to a substantially smaller list of possible candidates. Within the interval of interest for Q4 (Figure 6d), both *Mothers against dpp (Mad)* and α/β hydrolase 2 (*Hydr2)* have a *cis* eQTL in the King et al. ^72^ dataset. In addition, both of these genes are significantly differentially expressed between the high and low learning cohorts, with higher expression in the low learning cohort relative to the high learning cohort (*Mad:* log_2_FC: −0.37, p_adj_ = 0.015; *Hydr2*: log_2_FC= −0.38, p_adj_ = 0.015). While these genes are not significantly differentially expressed between the memory cohorts, the trend is in the same direction in the learning cohorts for both genes (*Mad:* log_2_FC: −0.13, p_adj_ = 0.67; *Hydr2*: log_2_FC= −0.19, p_adj_ = 0.45). *Mad* is a transcription factor that regulates the expression of BMP response target genes ^93^ and has also been shown to play a role in inhibiting synapse formation, as one part of two antagonistic signaling pathways that act together to produce the correct number of synapses ^94^. In the Q5 interval of interest (Figure 6e), just two genes are significantly differentially expressed, *Odorant-binding protein 28a (Obp28a)* and *Cyp4d21*, neither of which have evidence for a *cis* eQTL in the King et al. ^72^ dataset. Both of these are significant only between the learning cohorts, with higher expression in the high learning cohort (*Obp28a:* log_2_FC: 0.96, p_adj_ = 1.9 x 10^-4^; *Cyp4d21*: log_2_FC= 0.52, p_adj_ = 0.01). Neither of these genes have any previous evidence implicating their involvement in place learning or memory. There are six significantly differentially expressed genes within the Q9 interval of interest (Figure 6i), all of which have a *cis* eQTL in the King et al. ^72^ dataset. Of these, *mwh* is the only gene that is significantly differentially expressed between both the learning cohorts and the memory cohorts (Learning: log_2_FC: 0.37, p_adj_ = 0.004; Memory: log_2_FC= 0.36, p_adj_ = 0.01). In both cases, *mwh* is expressed more highly in the high performing cohorts. This gene is primarily involved in the formation of hairs ^73^. Appel et al.^95^ also found evidence for a role of the genes influencing mechanosensory bristles and electric shock avoidance in a different *D. melanogaster* mapping population, the DGRP. Within the Q11 interval (Figure 6k), there are two significantly differentially expressed genes, *Glutathione S transferase O1* (*GstO1*) and *ergic53*, both of which have a *cis* eQTL in the King et al. ^72^ dataset. *GstO1* is more highly expressed in the high learning cohort but is more lowly expressed in the high memory cohort, though the difference is only significant for learning (Learning: log_2_FC: 0.47, p_adj_ = 0.03; Memory: log_2_FC= −0.21, p_adj_ = 0.56). The gene *ergic53* is more highly expressed in the low learning and low memory cohorts relative to the high performing cohorts (Learning: log_2_FC: −0.41, p_adj_ = 0.03; Memory: log_2_FC= −0.37, p_adj_ = 0.11). An RNAi screen specific to neurons identified *ergic53* as a candidate gene involved in the perception of very high temperatures ^96^. Given the punishment in our learning paradigm is high temperature, this is a potentially exciting candidate gene; however, it is counterintuitive that the low learning cohort shows higher *ergic53* expression compared to the high learning cohort.

## Discussion

We successfully used a large multiparent population to identify multiple loci affecting place learning and memory in the fruit fly model system. We integrated an RNA-Seq dataset that provided a genome-wide characterization of differential expression between high and low performing cohorts to identify potential candidate genes at mapped loci. All of the identified loci represent novel loci not previously associated with place learning or memory.

Our study joins a small set of previous studies that have identified naturally occurring genetic variants influencing learning and/or memory ^24, 25, 46–50^, none of which examined place learning and memory. Selection and quantitative genetic approaches have long suggested that natural variants in the genome support variation in learning and memory. But specifically how genetic variation gives rise to better or worse performance has been challenging. Selection experiments in systems such as blow flies ^21^ and fruit flies ^22, 23^ have shown that natural genetic variants can be combined to yield better performance in learning tasks. However, only rarely have QTL for learning and memory been successfully mapped ^47–50^, as we have done here, and even more rarely have the specific genetic variants within a gene influencing learning and/or memory been identified (but see ^24, 25, 46^. As we build a larger collection of QTL that influence learning and memory in different systems, ideally validating the specific variants involved, we will gain a much greater understanding of the genetic mechanisms governing these processes.

Some of the major advantages of using a stable multiparent mapping panel such as the DSPR include the ability to measure multiple phenotypes on the same set of lines and the availability of additional datasets from other studies that can be integrated to address major questions. With these strengths we achieve a comprehensive picture of the genetic basis of both place learning and place memory. By measuring both phenotypes on the same set of lines, we showed that both traits are genetically correlated and identify specific loci that influence both traits. A constant challenge for QTL studies following mapping is determining which genes within the mapped interval are the most likely candidate genes. In traditional two-way QTL mapping, these intervals are typically very wide and usually include hundreds of genes ^97^. The multiple generations of crossing in the DSPR lead to smaller haplotype segment sizes and thus higher mapping resolution ^44^ however, single gene resolution is still not possible, requiring strategies for identification of likely candidate genes. Mapping the same QTL for multiple traits allowed us to focus on an even narrower interval of interest. This approach also allowed us to significantly narrow the interval of interest by considering only the area of overlap between the two BCI’s. By bringing in additional phenotyping, in the form of RNA-Seq of high performing and low performing cohorts, we could focus on genes that were significantly differentially expressed between these cohorts. Changes in gene expression in identified shared QTL in the same direction lent more support for a given gene. Finally, a particularly useful feature of the DSPR is the constantly increasing database of previous studies using this same set of lines (http://FlyRILs.org). We specifically used a large, genome-wide eQTL dataset ^72^ to identify which genes within our intervals of interest had previously been shown to harbor a *cis* eQTL. This is a powerful approach to identify genes that are affecting learning and memory via changes in gene expression. Other studies using multiparent populations have used a similar approach of data integration to more effectively identify candidate genes for traits such as resistance to toxins in fruit flies ^77, 98^, grain yield and flowering time in maize ^99^ and body weight in mice ^100^.

While we successfully identified potential candidate genes for many of our QTL, our approach does rely on several important assumptions. First, we focused on identifying candidate genes that are affecting learning and memory via regulatory variants, with a change in gene expression leading to a phenotypic change. However, it is also possible some of the genes underlying our mapped QTL are coding variants that do not lead to differences in the expression of the candidate gene. Our method of identifying candidate genes would miss such genes. Some of the few previously identified natural variants that have been found to influence learning and/or memory in other systems do show differences in gene expression ^24, 25^, and it has been hypothesized that many of the variants underlying QTL may be regulatory ^101, 102^. Second, we did not expose the lines we used for RNA-Seq to the heat box prior to sample collection. The training and memory test in the heat box happens over a short timescale, with the entire assay lasting just under 10 minutes, and thus we considered the differences in the baseline gene expression levels between cohorts to be a more likely predictor of the RILs’ performance on this test. Third, we performed RNA-Seq on pooled samples of female heads, which is an efficient way to average over a substantial amount of inter-individual and inter-RIL variability to pull out the set of genes consistently differentially expressed between high and low performers. However, it is also possible that this approach misses more subtle, but potentially important variation. For example, if differences in learning and memory are dependent on differences in gene expression in a specific set of cells, we could miss this effect using our approach. Future studies employing higher resolution expression analysis, such as the single cell approaches that are currently being used to characterize expression patterns across the fly brain ^103^, have the potential to provide this fine scale data. Finally, we note that we do not yet have additional functional information confirming that the genes we identify as candidate genes are causative, and it is certainly possible that the true causative gene for any QTL is one we have not identified here. Future follow up studies using approaches such as quantitative complementation ^104, 105^ or targeted studies of gene expression of potential candidate genes in the DSPR RILs along with learning and memory measurements will allow us to begin to determine exactly which of the genes in our intervals are causative.

Despite these caveats, our study represents a critical first step towards a more holistic characterization of the genetics of learning and memory in an important model system. The majority of previous investigations of the genetic basis of learning and memory have focused on single gene approaches. Several mutants have been identified that specifically affect place learning and/or memory ^80^. However, despite the success of these studies, a disconnect remains between the genes identified via single gene approaches and the identity of the genetic variants leading to individual-level variation in learning and memory in natural populations. In this study, none of the previously identified genes shown to specifically influence place learning and memory via mutant studies (e.g., *dunce, amnesiac, white, radish, rutabaga, arouser, and tribbles*) are within our QTL intervals. In addition, very few of these previously identified genes were significantly differentially expressed between high and low performing cohorts (see Results). This result is not wholly unexpected, given that learning and memory are expected to be highly complex, polygenic traits, and the mutations identified in single gene approaches are of large effect, are often quite deleterious, and would presumably be selected against in any natural population ^4, 42^. A similar disconnect between candidate gene approaches and the identity of natural causative variants has been found for other traits such as lifespan ^106–108^. However, in other cases, such as bristle number^109, 110^ and circadian rhythm^111, 112^ in *D. melanogaster,* QTLs have been identified that overlap with candidate genes identified via mutant analysis. Whether it is generally the case that a given gene identified as influencing a phenotype via mutant analysis typically also harbors more segregating variants influencing phenotypic variation will remain an open question until the identification of natural genetic variants underlying complex traits becomes more routine. Our study has taken a genome-wide approach, identifying loci involved in learning and memory without making major perturbations to the system, such as null mutations, blocking neurotransmitters, or eliminating major cell types. With this approach we have localized multiple QTL that influence whether a genotype learns and remembers well or poorly. Several of these QTL are likely pleiotropic loci, influencing both learning and memory. Others only map to a single phenotype and are potential loci contributing to some independence in these two processes. We also characterize genome-wide expression differences between high and low performing cohorts for these two phenotypes, providing a valuable profile of how high vs. low learning and high vs. low memory genotypes differ at the transcriptome level. Several other studies investigating the role of gene expression in learning have taken a before versus after approach, studying expression changes following training to identify which genes alter expression during and following learning ^113–115^. Our dataset instead provides a high performing versus low performing approach, identifying a set of hundreds of genes, whose regulatory differences potentially influence whether an individual will learn and remember well or poorly.

There is a growing appreciation for the complexity of learning and memory and the need to study these traits in realistic settings ^4, 116^. Recently there has been a shift from investigating single genes to identifying the sets of neurons and circuits involved in learning and memory ^54, 87, 117, 118^. In addition, mapping studies in humans are beginning to uncover the natural genetic variants influencing learning and memory ^25, 119^. Although it is still early, the challenges associated with studying human learning and memory suggest that a complete picture of how a genome can support fundamental learning and memory mechanisms remains in the future. Examination of these processes in animal models like fruit flies, with systems genetic resources such as the DSPR, provide a chance to identify mechanisms comprehensively through careful behavioral measures, the integration of phenotypic data across multiple levels of organization, and advanced quantitative genetics approaches.

## Acknowledgements

This project was only possible due to the leadership and enthusiasm of Dr. Troy Zars, who sadly did not live to see the work published. We thank Dr. Stuart Macdonald for supplying us with the DSPR RILs, Aditi Mishra for numerous helpful discussions about the project, and Research Computing Support Services at the University of Missouri, especially Christopher Bottoms and Jacob Gotberg, for their bioinformatics computing support. Kevin Middleton provided helpful comments that improved the manuscript and produced the fly image used in Figure 1. This material is based upon work supported by the National Science Foundation under Grant No. IOS-1654866 (T.Z. and E.G.K.), National Institutes of Health R01GM117135 (E.G.K.), MU Research Council Grant URC-16-007 (T.Z.), National Institute of Health IMSD R25GM056901 (P.A.W-S.), and an HHMI Gilliam Fellowship (P.A.W-S.).

