## Supporting Information for "Multiple genetic loci affect place learning and memory performance in *Drosophila melanogaster*"

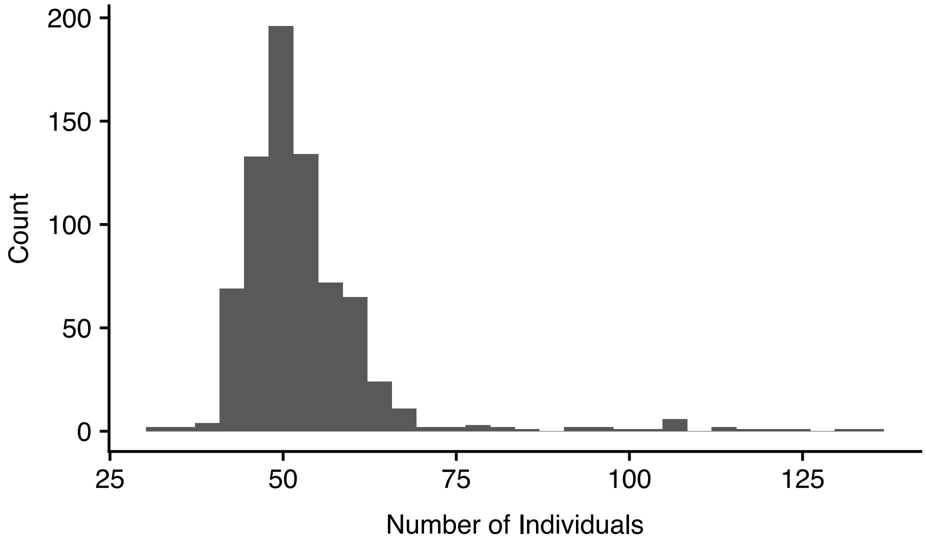


Figure S1: Histogram of the number of individuals assayed per DSPR RIL. The minimum number of individuals assayed was 31, with an average of 53.

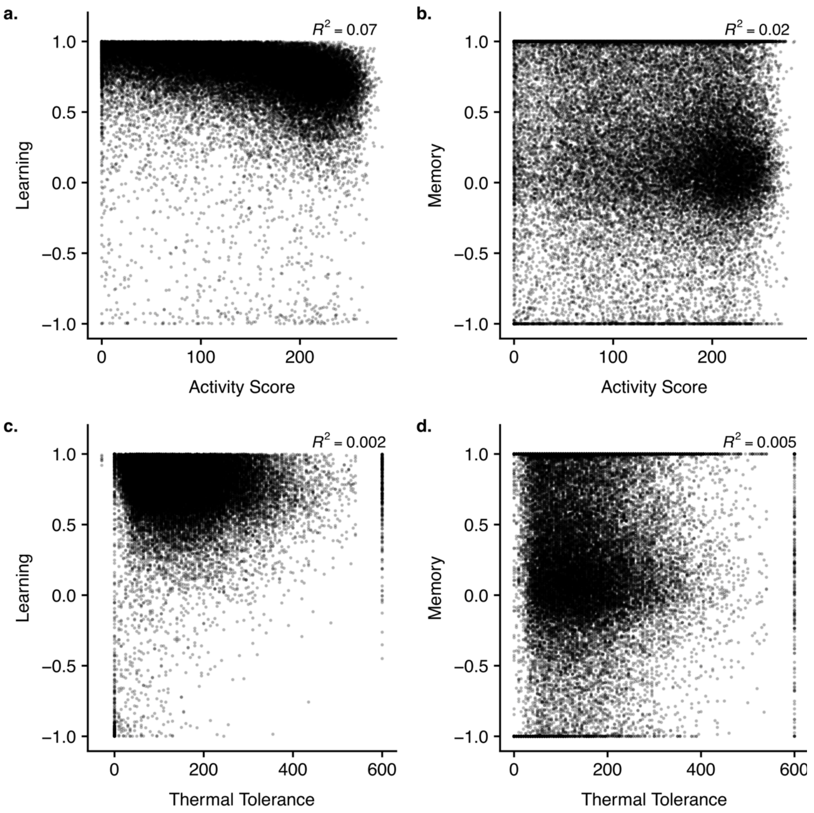


Figure S2: a & b) The relationship between individual activity and learning (a) and memory (b). c & d) The relationship between thermal tolerance and learning (c) and memory (d). The percent variance explained ($R^2$) is shown in the top right corner of each plot. Points are semi-transparent such that overlapping points will appear darker.

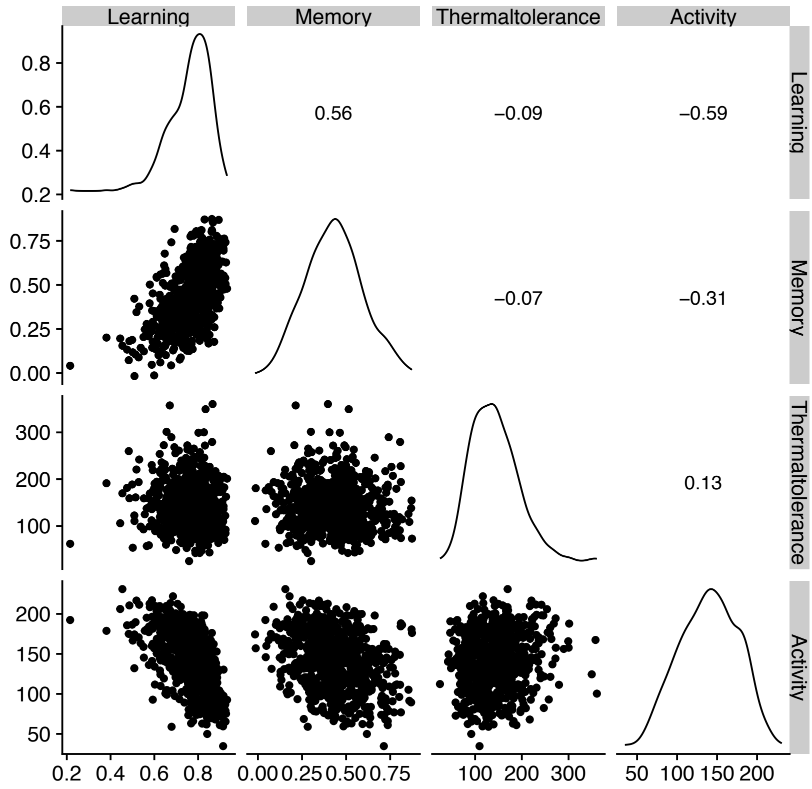


Figure S3: Phenotypic patterns of the average scores for each RIL for learning, memory, thermal tolerance, and activity. Density plots for each phenotype are along the diagonal. In the bottom triangle, pairwise plots showing the relationships between each pair of traits are shown. In the upper triangle, the value of the Pearson's correlation coefficient is shown.

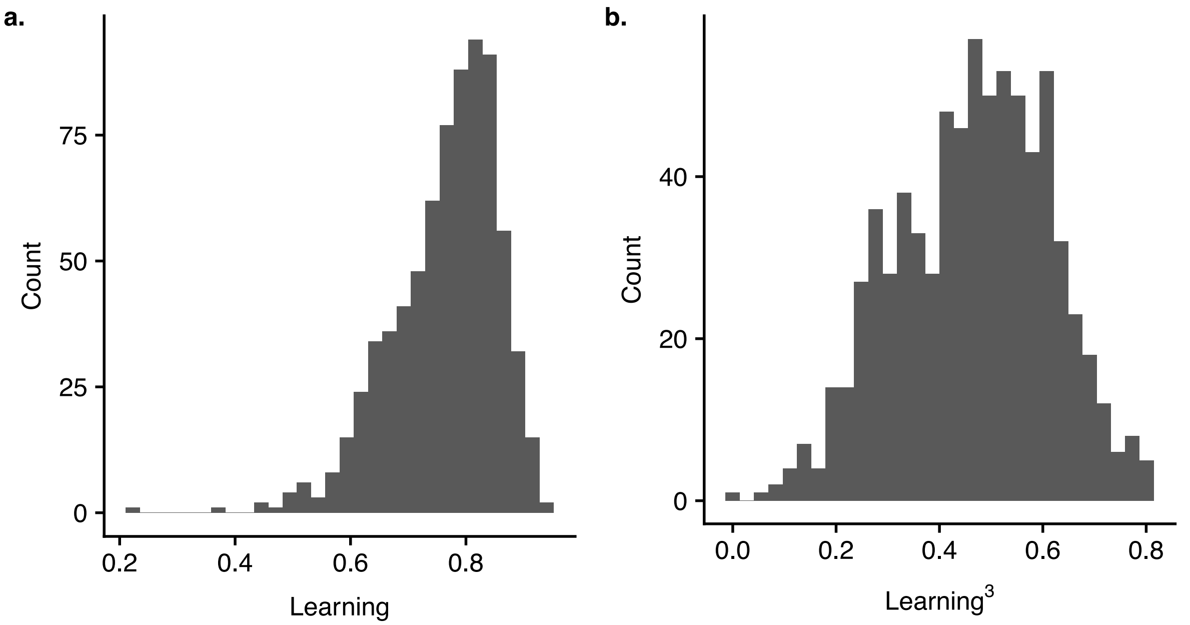


Figure S4: Histograms of the average learning scores for 743 RILs. The untransformed average scores are shown in (a) and the power transformed average scores are shown in (b).

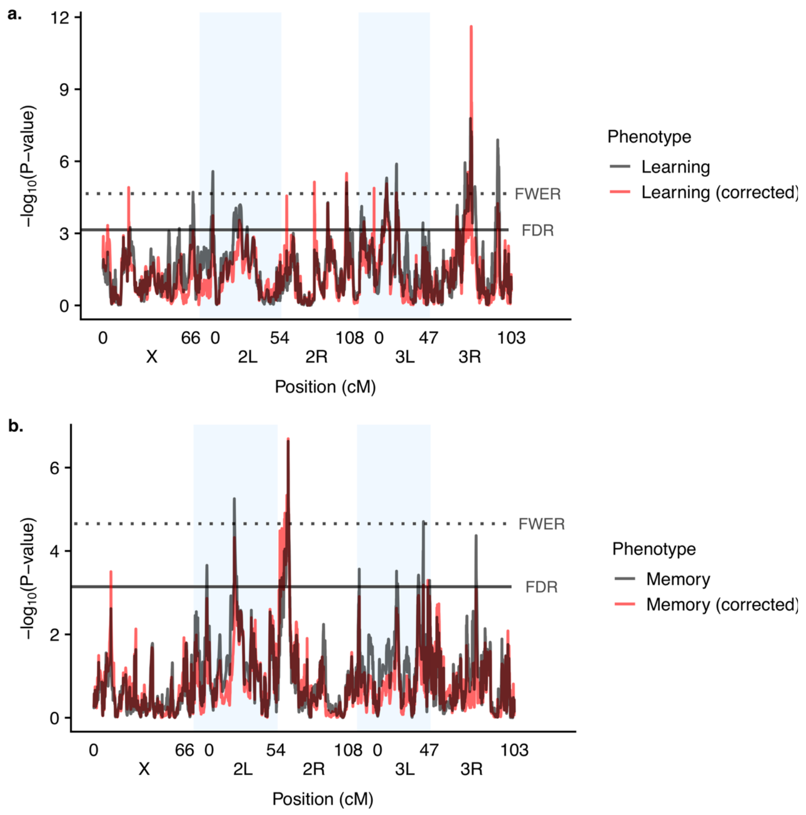


Figure S5: *Genome scan of learning (a) and memory (b) phenotypes.* Different colors denote the genome scan for raw learning and memory scores versus scores statistically corrected for individual activity levels. Dotted horizontal line denotes the 5% family-wise error rate threshold and the solid horizontal line denotes the 5% false discovery rate. Shaded blue boxes denote chromosome arms.

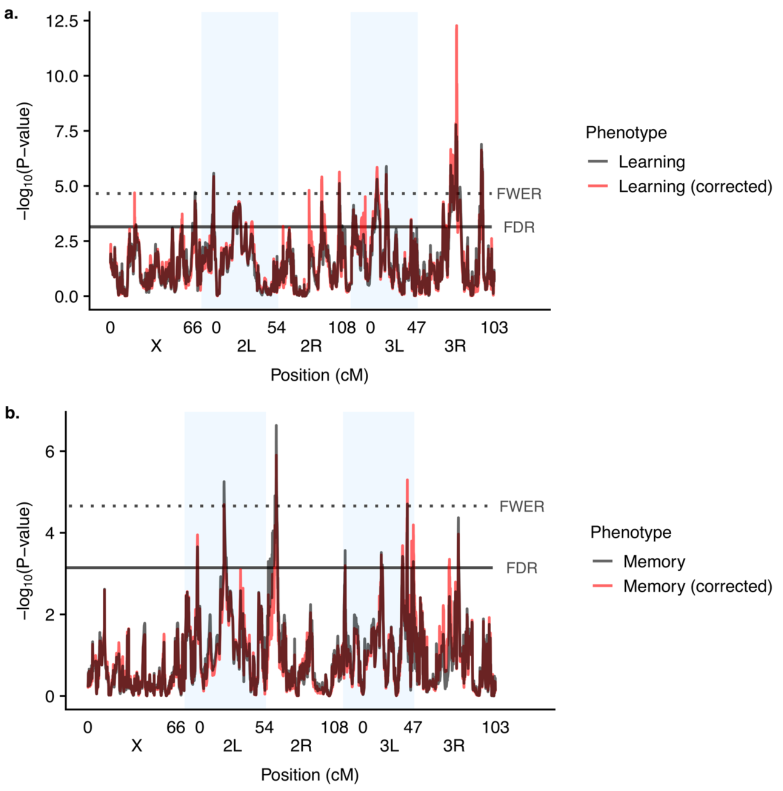


Figure S6: *Genome scan of learning (a) and memory (b) phenotypes.* Different colors denote the genome scan for raw learning and memory scores versus scores statistically corrected for individual thermal tolerance levels. Dotted horizontal line denotes the 5% family-wise error rate threshold and the solid horizontal line denotes the 5% false discovery rate. Shaded blue boxes denote chromosome arms.
